# Harmine enhances the activity of the HIV-1 latency-reversing agents ingenol A and SAHA

**DOI:** 10.1101/2020.01.31.927368

**Authors:** Jared P. Taylor, Lucas H. Armitage, Daniel L. Aldridge, Melanie N. Cash, Mark A. Wallet

## Abstract

Infection of HIV-1 remains incurable because long-lived, latently-infected cells persist during prolonged antiretroviral therapy. Attempts to pharmacologically reactivate and purge the latent reservoir with latency reactivating agents (LRAs) such as protein kinase C (PKC) agonists (e.g. ingenol A) or histone deacetylase (HDAC) inhibitors (e.g. SAHA) have shown promising but incomplete efficacy. Using the J-Lat T cell model of HIV latency, we found that the plant-derived compound harmine enhanced the efficacy of existing PKC agonist LRAs in reactivating latently-infected cells. Treatment with harmine increased not only the number of reactivated cells but also increased HIV transcription and protein expression on a per-cell basis. Importantly, we observed an additive effect when harmine was used in combination with ingenol A and the HDAC inhibitor SAHA. An investigation into the mechanism revealed that harmine, when used with LRAs, increased the availability of transcription factors needed for viral reactivation such as NFκB, MAPK p38, and ERK1/2. We also found that harmine treatment resulted in reduced expression of HEXIM1, a negative regulator of transcriptional elongation. Despite harmine’s reported inhibitory effects on DYRK1A and consequent enhancement of NFAT signaling, the HIV reactivating effects of harmine occurred independent of DYRK1A and NFAT. Harmine increases the efficacy of LRAs by increasing the availability of HIV-1 transcription factors and decreasing expression of HEXIM1. Combination therapies with harmine and LRAs could benefit patients by achieving deeper reactivation of the latent pool of HIV provirus.

## Introduction

Infection with human immunodeficiency virus 1 (HIV) remains incurable because of the ability of this virus to permanently integrate its genetic material into the host genome of infected cells. The ssRNA genome of HIV requires a stable intermediate in the form of an integrated proviral DNA to complete its life cycle. This integrated DNA is subject to the same regulatory mechanisms as host genes including requirements for active transcription factors as well as epigenetic control of chromatin structure and accessibility ^1^.

Like host genes, the HIV genome may enter a state of non-expression wherein viral mRNA and proteins are not expressed ^2^. Whether this non-expression is through neglect (lack of stimuli required to drive the HIV LTR promoter) or active suppression (binding of inhibitory proteins to the LTR or epigenetic silencing of the locus), the outcome is the same – HIV is not produced, and the virus remains hidden from host immunity. In this state, HIV endures for as long as the infected cell and all progeny from future cell divisions. If replication-competent proviral genomes are harbored, an individual is at risk for viral reactivation.

Even after HIV replication has been pharmacologically suppressed so that the virus is undetectable in peripheral blood, cessation of treatment leads to the resumption of the HIV/AIDS clinical progression ^3^. Elimination of the latent reservoir or significant reduction of the size of the reservoir are seen as the only real hopes for a sterilizing or functional cure, respectively. A large body of work is focused on pharmacological strategies to “purge” the latent reservoir. The most widely studied approach is known as “shock and kill” or “kick and kill” ^4,5^. The goal is to simultaneously reactivate all (or most) replication-competent latent HIV while maintaining antiretroviral therapy (ART). Ideally, upon re-expression of viral mRNA and proteins, the reservoir cells will die through either cytopathic effects of the virus or through immune mechanisms (e.g. cytotoxic T lymphocytes [CTL] or natural killer [NK] cells). What has been lacking is a safe and effective pharmacological method to potently reactivate latent HIV *in vivo.*

HIV latency is predominantly controlled by two basic mechanisms – chromatin accessibility and transcription factor expression/activation/localization. Epigenetic regulation of the HIV integration site through modification of histone proteins by histone deacetylase (HDAC) enzymes effectively silences HIV mRNA expression. HDAC inhibitors such as vorinostat (SAHA) can elicit HIV replication from latently infected cells *in vitro* ^6–9^. *In vivo*, vorinostat can induce some reactivation and increase plasma HIV RNA in subjects receiving ART ^10^. However, thus far HDAC inhibitors have not been able to significantly reduce the pool of latently infected cells.

Transcription of HIV mRNA relies on the interaction of the viral protein Tat and the pTEFb complex. The pTEFb complex is a positive regulator of transcription elongation and the interaction of Tat with pTEFb is required for efficient elongation of HIV transcripts ^11,12^. The activation of host transcription factors such as NFκB and MAPK are also required for reactivation of HIV from latency. These transcription factors are normally sequestered in the cytoplasm as inactive proteins. Numerous early studies of HIV latency have focused on treatments that stimulate T cell activation including cytokines (IL-2 and IL-7), ligation of surface proteins (PHA, anti-CD3) or chemical stimulators of signaling pathways such as protein kinase C (PKC) agonists. NFκB, in particular, is a potent inducer of HIV mRNA expression and several PKC agonists have been validated for their ability to elicit reactivation of latent HIV. These treatments are even more potent when paired with complementary drugs that target HDACs or BRD4. Other transcription factors such as SP1, STATs, and IRFs play roles in regulating the HIV LTR. The NFAT family of transcription factors may also regulate HIV latency since the HIV LTR contains NFAT transcription factor binding sites and NFAT can enhance HIV mRNA expression. NFAT, like NFκB, is sequestered as an inactive protein in the cytoplasm unless activated by specific upstream signals. However, unlike NFκB, NFAT is hyperphosphorylated in its inactive state ^13^. This hyperphosphorylation is driven by a dual-specificity kinase DYRK1A ^14,15^ and NFAT only becomes de-phosphorylated following calcium-dependent calmodulin signaling ^13^.

Harmine is a naturally-occurring tricyclic β-carboline alkaloid with hallucinogenic properties that is derived from the plant *Banisteriopsis caapi* as well as others ^16^. Harmine has been shown to be an inhibitor of DYRK1A ^17–19^. Harmine’s primary target is monoamine oxidase A (MAO-A)^20–23^, but a well-reported role for inhibition of DYRK1A leading to enhanced NFAT activity has been reported ^17–19,24^. The crystal structure of harmine complexed with DYRK1A has been solved confirming its binding to the ATP-binding pocket of DYRK1A ^25^. Because of its ability to augment NFAT signaling, we undertook a study of harmine and other DYRK1A inhibitors to determine if these compounds could enhance HIV reactivation alone or in combination with other known latency-reversing agents (LRAs). We hypothesized that DYRK1A inhibitors would augment reactivation with LRAs through increased NFAT availability. Here we report that harmine and another DYRK1A inhibitor, INDY, boost HIV reactivation by PKC agonists. Interestingly, the effect was independent of NFAT activity; harmine was effective at boosting LRAs even when no evidence of NFAT activity was detected. In addition, CRISPR knockout of DYRK1A did not mimic the effects of harmine or modulate the efficacy of harmine for boosting HIV reactivation, Instead, we found that harmine enhances MAPK and NFκB signaling leading to increased HIV transcription. Using whole-genome microarray we observed that harmine modulates expression of key pTEFb components. *HEXIM1* expression is significantly downregulated by harmine whereas the cyclin *CCNT2* was upregulated. We conclude that harmine regulates pTEFb complex heterogeneity and establishes an environment that is more conducive to HIV reactivation. Combination treatments with harmine and LRAs may prove efficacious *in vivo*.

## Results

### DYRK1A inhibitors enhance the efficacy of T cell activating PKC agonists

PKC agonists have been previously shown to reactivate latent HIV^26–33^. We expected reactivation with PKC agonists to be enhanced by combinatorial treatment with DYRK1A inhibitors, harmine or INDY (**Supplemental Figure 1A**). To test this, we utilized J-Lat 5A8 reporter cells, which are latently infected with a full-length provirus that expresses GFP in place of Nef as an indicator of LTR activity. J-Lat 5A8 cells were pretreated with harmine or INDY and reactivated with the PKC agonist ingenol A. Both harmine and INDY failed to cause any measurable HIV reactivation when used alone, however, harmine and INDY boosted the reactivation effects of ingenol A (**Figure 1A**). This effect could be seen with doses as low as 5 µM in J-Lat cells and as low as 2.5 µM in primary CD4+ T cells (**Supplemental Figure 1B&C**). Toxicity caused by harmine and INDY was seen with doses above 25 µM (**Supplemental Figure 1D**), therefore, a dose of 20 µM was used for experiments.

**Figure 1.**
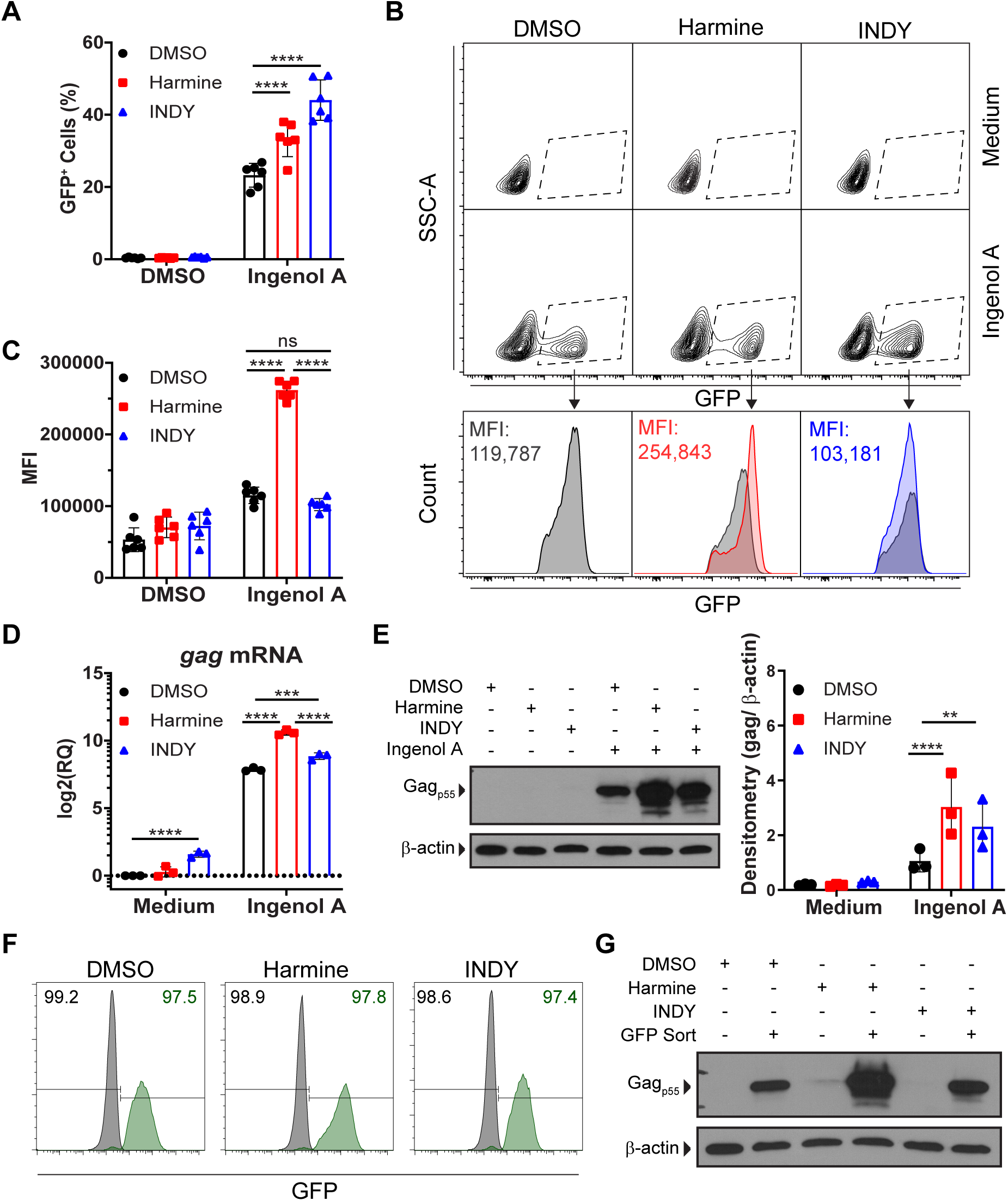
DYRK1A inhibitors enhance the efficacy of PKC agonist ingenol A. **(A)** The percentage of GFP^+^ J-Lat 5A8 cells after reactivation with ingenol A in the presence of inhibitors (n = 6), **(B)** representative flow cytometry plots of the mean fluorescence intensity of GFP^+^ cells, and **(C)** summary mean fluorescence intensity data (n = 6). **(D)** *gag* mRNA expression determined by RT-qPCR. **(E)** Gag protein expression determined by western blot with representative blot and densitometry (n = 3). GFP^+^ and GFP^-^ cells were sorted after reactivation with ingenol A. **(F)** Post-sort purity measured by flow cytometry. The percentage of GFP^-^ population (black) and the GFP^+^ population (green). **(G)** Gag protein expression of post-sort GFP^+^ cells measured by western blot. Error bars represent standard deviation. Statistical analysis was performed by two-way ANOVA and corrected for multiple comparisons by Tukey’s test. **p<0.01; ***p<0.001; ****p<0.0001; ns = not significant.

When analyzing flow cytometry data, we recognized a unique phenomenon in the J-Lat reactivation experiments. Not only was harmine increasing the frequency of cells that became GFP^+^, but it also appeared that the brightness of each GFP^+^ cell was increased when harmine was included (**Figure 1B-C**). J-Lat cells were then activated with increasing doses of ingenol A, PMA, or TNF in the presence of DMSO, harmine, or INDY. Both harmine and INDY boosted the frequency of ingenol A or PMA-activated cells whereas only INDY had a positive effect on reactivation with TNF treatment (**Supplemental Figure 2**). When gating on only the GFP^+^ cells, it became clear that harmine increases the amount of GFP expressed in each reactivated cell (**Supplemental Figure 2**). This finding suggests that harmine works through a mechanism that is unique from INDY. It also seems to indicate that the boosting effect of harmine is working through at least two biological pathways – one that increases the sensitivity of cells to activating stimuli (reduced dose of stimulus required to activate HIV) and a second pathway that enhances the magnitude of HIV LTR activity in cells where reactivation occurs.

**Figure 2.**
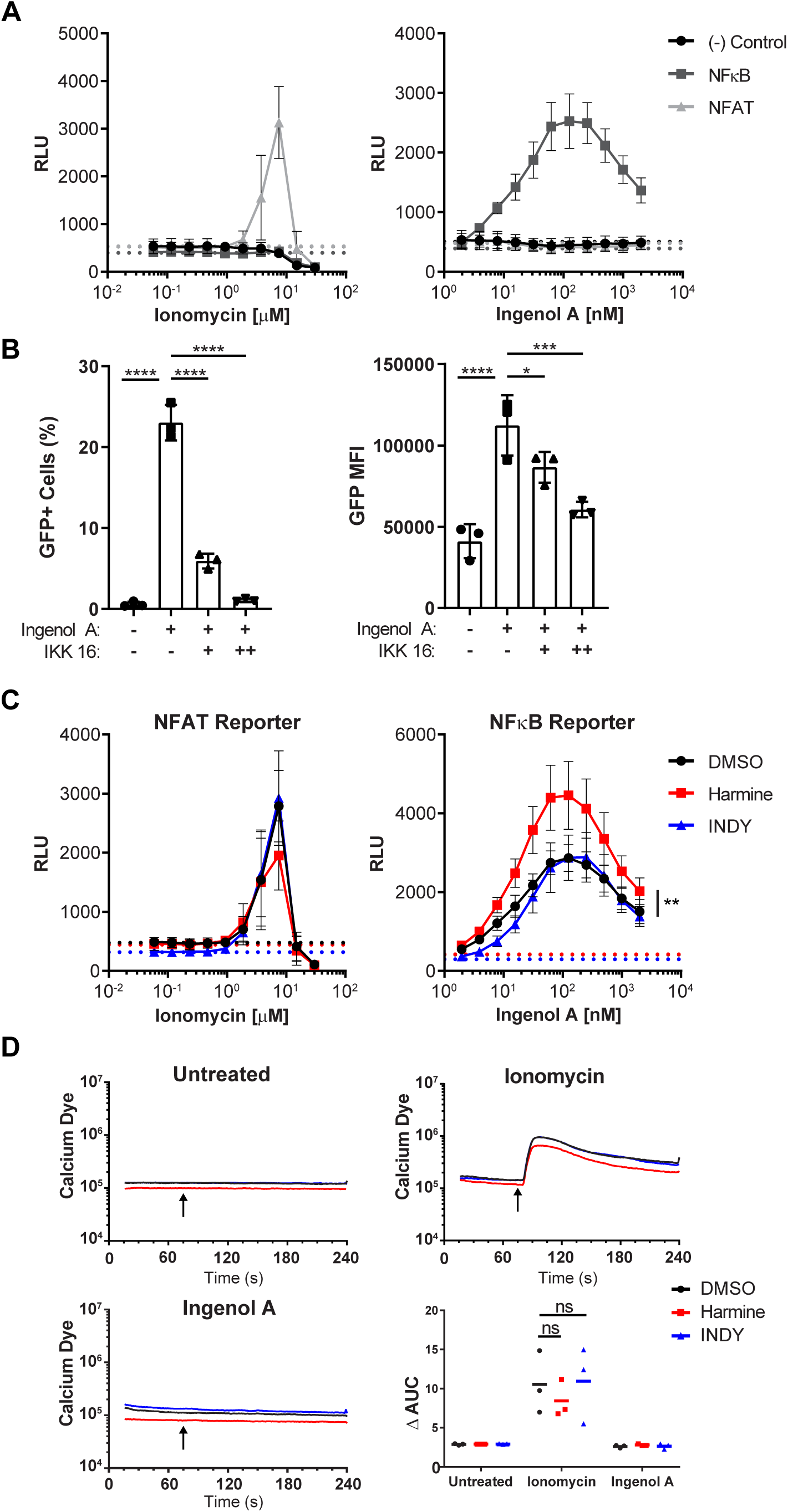
Harmine’s boosting effect is through NFκB, not NFAT. **(A)** Ionomycin or ingenol A were titrated on J-Lat 5A8 luciferase reporter cells. The dashed lines indicate the luciferase activity with no ionomycin or ingenol A treatment. **(B)** NFAT or NFκB luciferase reporter cells reactivated by ionomycin or ingenol A in the presence of inhibitors. Dashed lines represent luciferase activity with no ionomycin or ingenol A treatment. **(C)** J-Lat 5A8 cells were stained with Calcium Sensor Dye eFluor 514 (2 µM) and treated with DMSO, harmine, or INDY for 30 minutes. Geometric mean fluorescence was measured by flow cytometry for 240 seconds. Ionomycin (250 nM) or ingenol A (50 nM) was added after 75 seconds (arrows). Plots show mean geometric fluorescence intensity over time and the change in area under the curve (n = 3). **(D)** J-Lat 5A8 cells were pretreated with DMSO or an IκB kinase inhibitor, IKK 16, at 1 µM (+) or 10 µM (++) followed by ingenol A treatment (31.25 nM) for 18 hours (n = 3). Error bars represent standard deviation. Statistical analysis was performed by one-way-ANOVA corrected for multiple comparisons with Dunnett’s test (D) and one-way-ANOVA of the area under the curve corrected for multiple comparisons by Tukey’s test (C). *p<0.05; ***p<0.001; ****p<0.0001; ns = not significant; RLU = relative light units.

For a more direct measure of HIV-encoded genes/proteins, J-Lat cells were activated with ingenol A in the presence or absence of harmine or INDY. Here, like GFP, *gag* mRNA expression was induced by ingenol A + DMSO and the expression was markedly increased with the addition of harmine (**Figure 1D**). Harmine alone failed to induce *gag* expression. However, INDY alone did result in increased gag expression (**Figure 1D**) even though INDY treatment alone did not result in increased activation of J-Lat 5A8 cells (**Figure 1A**). Next, western blot analysis was performed to measure gag protein expression. Gag protein expression was induced by ingenol A alone and markedly increased with the addition of harmine or INDY. The effect was stronger for harmine than with INDY (**Figure 1E**). To better measure the amount of gag being expressed on a per cell basis, flow cytometry sorting was used to separate GFP^+^ vs GFP^-^ cells after ingenol A treatment in the presence or absence of harmine or INDY. Gag protein expression was measured in the GFP^+^ and GFP^-^ cells by western blot. The GFP^+^ cells that were treated with ingenol A + harmine expressed more gag protein than GFP^+^ cells that were treated with ingenol A + INDY (**Figure 1 F&G**). Again, these findings indicate that harmine is mediating two distinct effects on HIV latency. One effect increases the sensitivity of cells to activating LRAs whereas a second effect increases the expression level of HIV genes/proteins. This finding leads to two key questions: 1) Does harmine enhance HIV reactivation through enhanced NFAT activity? and 2) Is DYRK1A the target of harmine in HIV latency models?

### Harmine boosts HIV reactivation independent of NFAT

DYRK1A negatively regulates NFAT by hyperphosphorylation, which excludes NFAT from the nucleus. Only through calcium-dependent calmodulin activation is NFAT de-phosphorylated resulting in nuclear translocation and activation of NFAT dependent genes such as IL-2. The HIV LTR contains at least two NFAT binding motifs and since harmine is a known inhibitor of DYRK1A, we wanted to determine if the anti-latency effects of harmine are partly due to enhanced NFAT activity.

First, we wanted to determine if harmine affects NFAT activity in T cells. Jurkat T cells were transduced with lentiviral luciferase reporter constructs for NFAT, NFκB or a negative control virus with luciferase gene but no promoter. When reporter cells were treated with ionomycin alone – a well-known activator of calmodulin and NFAT – only the NFAT reporter cells expressed luciferase (**Figure 2A**). When reporter cells were treated with ingenol A alone, only the NFκB cells expressed luciferase (**Figure 2A**). This suggests that reactivation of J-Lat 5A8 cells is dependent on NFκB, not NFAT. Furthermore, treatment of J-Lat 5A8 cells with IKK16, an inhibitor of NFκB signaling, significantly reduced the percentage of reactivated cells as well as the MFI of GFP^+^ cells in a dose-dependent manner (**Figure 2B**). To determine if harmine was influencing NFAT or NFκB activity, NFAT and NFκB cells were treated with ingenol A with harmine or INDY. Harmine had no effect on ionomycin-induced luciferase in the NFAT reporter cells. However, harmine significantly boosted luciferase activity in the NFκB reporter cells compared to INDY. INDY had no effect on luciferase expression in any of the cell lines (**Figure 2C**). These data suggest that the boosting effects of harmine and INDY are through different mechanisms.

Second, we wanted to determine if harmine or INDY affected calcium flux upstream of NFAT activation. Jurkat cells were stained with a calcium-sensitive dye and treated with ionomycin or ingenol A. Ionomycin induced a calcium flux as expected but ingenol A did not. Neither harmine nor INDY had any effect on calcium flux regardless of agonist treatment (**Figure 2D**). Harmine is primarily affecting the NFκB pathway with no significant involvement of NFAT or the calcium flux. Thus, harmine and INDY are working through different mechanisms. Harmine appears to be acting upon the NFκB pathway.

We next wanted to determine if harmine was only affecting the NFκB pathway or if it was also affecting other pathways downstream of PKC. J-Lat cells cultured with PMA +/- harmine were analyzed for phospho-ERK1/2, phospho-MAPKp38, and phospho-AKT levels. We found that harmine markedly boosted phospho-ERK1/2 and phospho-MAPKp38 levels induced by PMA (**Figure 3A**). We found no difference in levels of phospho-AKT (**Figure 3A**). Inclusion of an ERK inhibitor, U0126, resulted in dose-dependent suppression of ingenol A-induced HIV reactivation (**Figure 3B**). Importantly, while the inhibitor markedly reduced the frequency of GFP^+^ cells, it also reduced the MFI of GFP^+^ cells (**Figure 3C**). This reduction in the MFI of the GFP^+^ cells was more pronounced than the reduction in MFI seen with the inhibition of NFκB. Thus, it appears that harmine boosts the frequency of GFP^+^ cells by enhancing sensitivity to PKC agonists as well as increases the magnitude of LTR activity independent of a strong activating signal.

**Figure 3.**
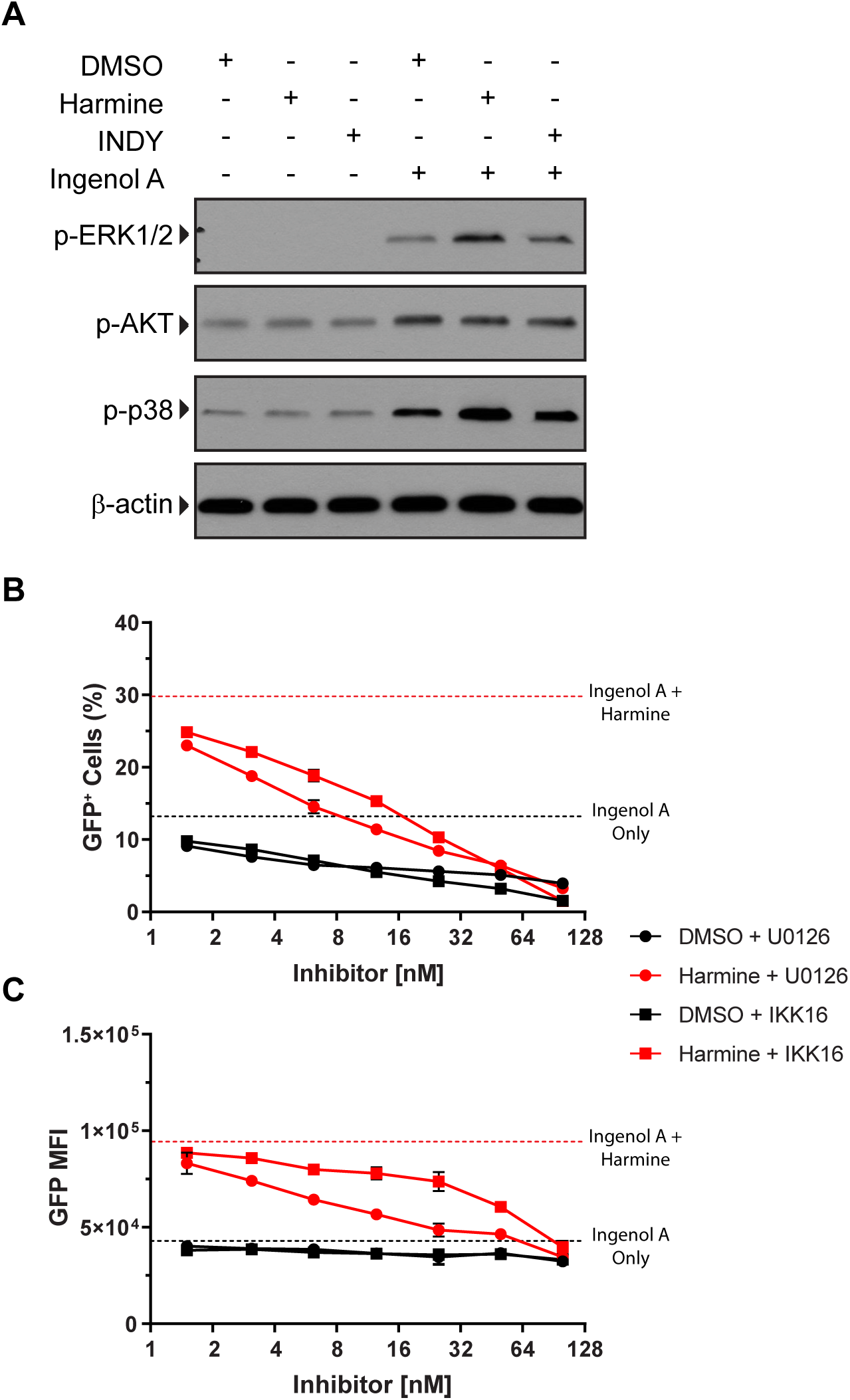
Harmine boosts phospho-ERK1/2 and phospho-p38 levels after ingenol A stimulation. J-Lat 5A8 cells were reactivated with ingenol A (31.25 nM) in the presence of inhibitors. **(A)** Whole cell lysates were analyzed by western blot for phospho-ERK1/2, phospho-AKT, and phospho-p38. **(B)** J-Lat 5A8 cells were pretreated with DMSO, a MEK inhibitor (U0126), or an IκB kinase inhibitor (IKK 16) for 30 minutes followed by overnight treatment with ingenol A (100 nM). GFP expression was assessed by flow cytometry. The percentage and **(C)** mean fluorescence intensity of the GFP^+^ cells are shown. Dash lines represent treatment with ingenol A alone.

### DYRK1A expression is not required for harmine’s positive effects on HIV reactivation

Since we observed that the boosting effect of harmine was independent of NFAT activity and altered MAPK signaling, we wanted to confirm that harmine specifically inhibited DYRK1A. A J-Lat cell line lacking DYRK1A expression was derived by CRISPR/Cas9 gene targeting (**Figure 4A**). Because harmine is an inhibitor of DYRK1A, we anticipated that J-Lat cells lacking DYRK1A would behave similarly to harmine treated cells with increased GFP expression in response to activating LRAs. Surprisingly, DYRK1A expression was not required for harmine or INDY to increase the frequency of GFP^+^ cells or for harmine to increase the MFI of GFP^+^ cells (**Figure 4B**). These data show that harmine acts through a different pathway altogether and does not require DYRK1A for boosting HIV reactivation with LRAs.

**Figure 4.**
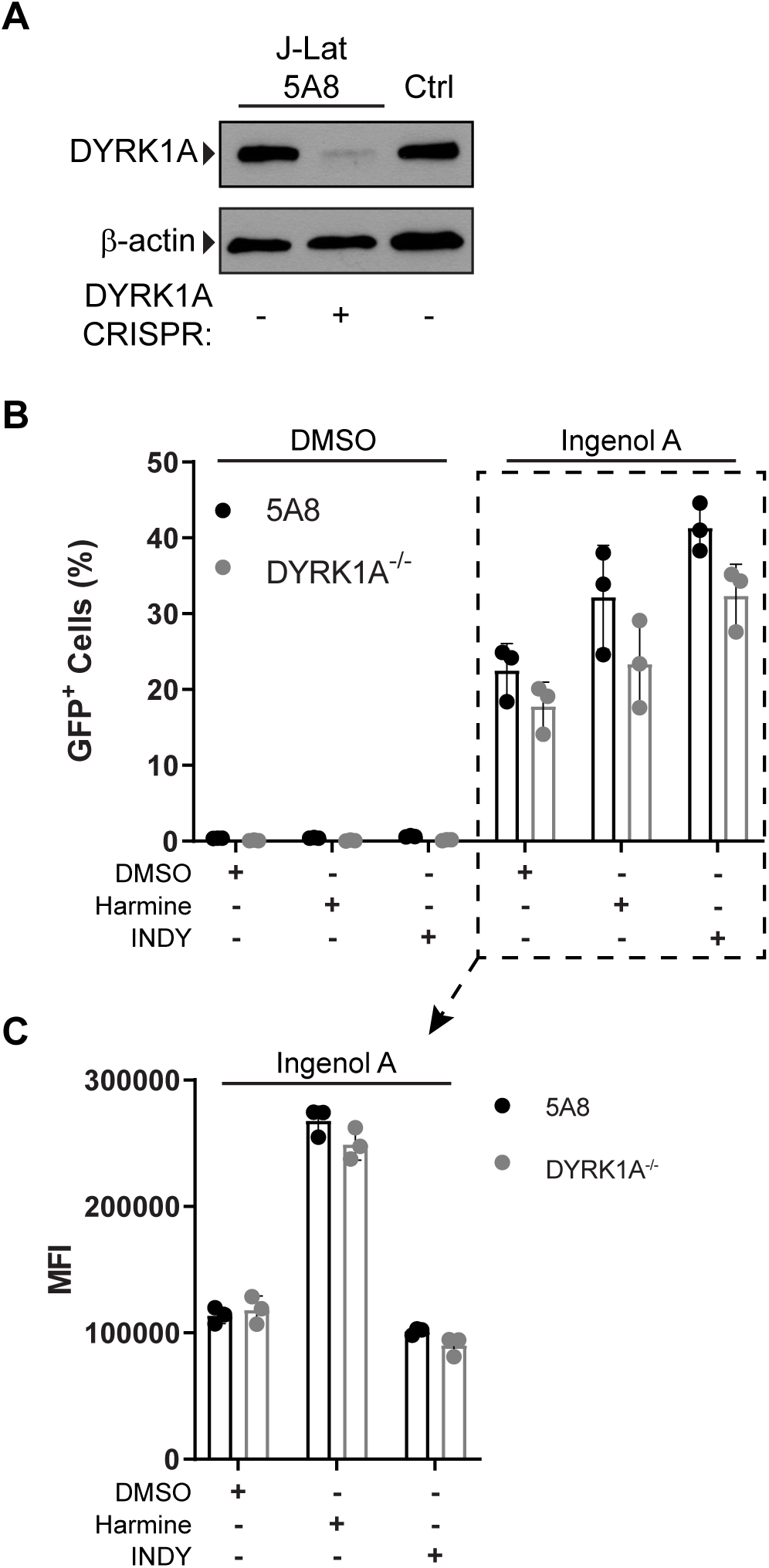
Harmine boosts independently of DYRK1A. DYRK1A was knocked out with CRISPR/Cas9 in J-Lat 5A8 cells. **(A)** Western blot of J-Lat 5A8 cells after treatment with CRISPR compared to Jurkat cells (Ctrl). The CRISPR knockout cells were reactivated with ingenol A (31.25 nM) in the presence of inhibitors. **(B)** The percentage of GFP^+^ cells and the **(C)** mean fluorescence intensity of GFP^+^ cells was measured by flow cytometry. Error bars represent standard deviation.

### Harmine downregulates HEXIM1 expression

To better understand how harmine is modulating HIV reactivation, we utilized whole-genome microarray to determine which genes were differentially expressed in cells treated with harmine. J-Lat cells were treated with DMSO, harmine, PMA, or harmine + PMA. Microarray analysis showed that there were 35 transcripts that were significantly upregulated or downregulated by at least 2-fold between PMA and PMA + harmine treatments. A complete list of the 35 significantly upregulated or downregulated genes are presented in **Supplemental Table 1.** Fourteen of these transcripts correspond to coding genes (**Figure 5A and Table 1**) and the remaining 21 transcripts were non-coding transcripts. The gene that was upregulated the most in PMA + harmine treatment compared to PMA treatment alone was *CCNT2*, which codes for the cyclin T2 protein. The gene that was downregulated the most in PMA + harmine treatment compared to PMA treatment alone was *HEXIM1.* Both genes code for proteins that play a role in transcriptional elongation ^34^. The HIV-encoded transcription factor Tat competes with HEXIM1 for binding to the pTEFb complex to promote HIV transcription ^35^. Downregulation of *HEXIM1* would thus result in less negative regulation of the pTEFb complex resulting in more Tat binding to pTEFb promoting transcript elongation. Similarly, upregulation of cyclin T2 would be expected to increase elongation of transcripts. This combination of effects on transcription explains why harmine treatment not only increases the percentage of reactivated cells but also increases the number of viral transcripts on a per-cell basis.

**Table 1.** PMA-Induced Gene Expression with Harmine.

| **Gene Symbol** | **Fold-Change**<br>**(PMA + harmine vs. PMA alone)** |
| --- | --- |
| CCNT2 | 2.2142 |
| SOX30 | 2.10846 |
| TMEM2 | -2.0027 |
| RAB23 | -2.03377 |
| ANKRD50 | -2.12436 |
| IL27RA | -2.1761 |
| RASA2 | -2.3009 |
| SLC7A11 | -2.37159 |
| LRRC8B | -2.3876 |
| SKIL | -2.4415 |
| USP27X | -2.44667 |
| CHAC1 | -2.52968 |
| HEXIM1 | -3.29431 |

**Figure 5.**
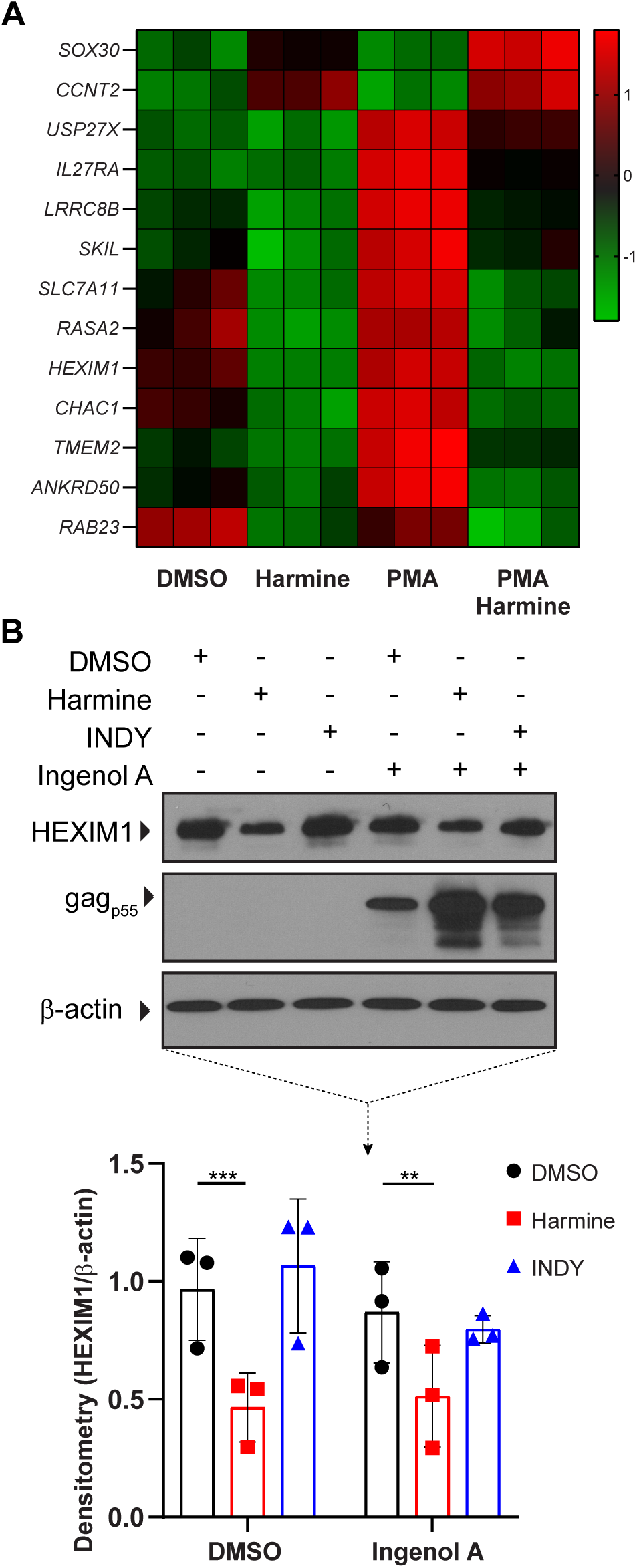
Harmine downregulates HEXIM1 expression. J-Lat 5A8 cells were treated pretreated with DMSO, harmine, or INDY for 30 minutes followed by PMA stimulation for two hours and analyzed by whole transcriptome microarray. **(A)** Hierarchical clustering of transcripts that were upregulated or downregulated by at least two-fold between PMA and harmine + PMA treatments. **(B)** Representative western blot analysis of HEXIM1 expression and densitometry (n = 3). Error bars represent standard deviation. Statistical analysis was performed by two-way-ANOVA corrected for multiple comparisons with Tukey’s test. **p<0.01; ***p<0.001.

To confirm the microarray findings, we treated J-Lat cells with ingenol A in the presence or absence of harmine or INDY. Western blot analysis showed that harmine treatment alone or with ingenol A leads to a significant reduction in HEXIM1 protein expression, whereas INDY treatment has no effect (**Figure 5B**). The decrease in HEXIM1 protein expression corresponded to an increase in the expression of gag protein. Thus, harmine treatment alone downregulates HEXIM1 and boosts the efficacy of subsequent LRA stimulation.

### Harmine boosts the efficacy of SAHA-induced HIV reactivation

We wanted to determine if harmine also enhanced other LRAs independent of the PKC pathway. We observed that harmine also boosts the efficacy of SAHA (vorinostat), an HDAC inhibitor, to enhance viral reactivation. Harmine only boosts the MFI of SAHA-reactivated cells and not the frequency of reactivated cells (**Supplemental Figure 3**) because SAHA reactivation is independent of NFκB or NFAT pathways. We next tested the combinatorial effects of ingenol A, SAHA, and harmine on HIV reactivation. SAHA and harmine both boosted the percentage of ingenol A-reactivated cells and that the combination of SAHA and harmine had a more potent effect on ingenol A-induced reactivation (**Figure 6A**). SAHA did not boost the MFI of ingenol A-reactivated GFP^+^ cells. However, harmine did boost the MFI of ingenol A-reactivated GFP^+^ cells and the combination of harmine and SAHA had an even greater boosting effect (**Figure 6B**). Similarly, we found that while harmine + ingenol A increases *gag* mRNA (**Figure 6C**) and gag protein (**Figure 6D**) expression compared to ingenol A alone that harmine + SAHA + ingenol A induced significantly more *gag* expression.

**Figure 6:**
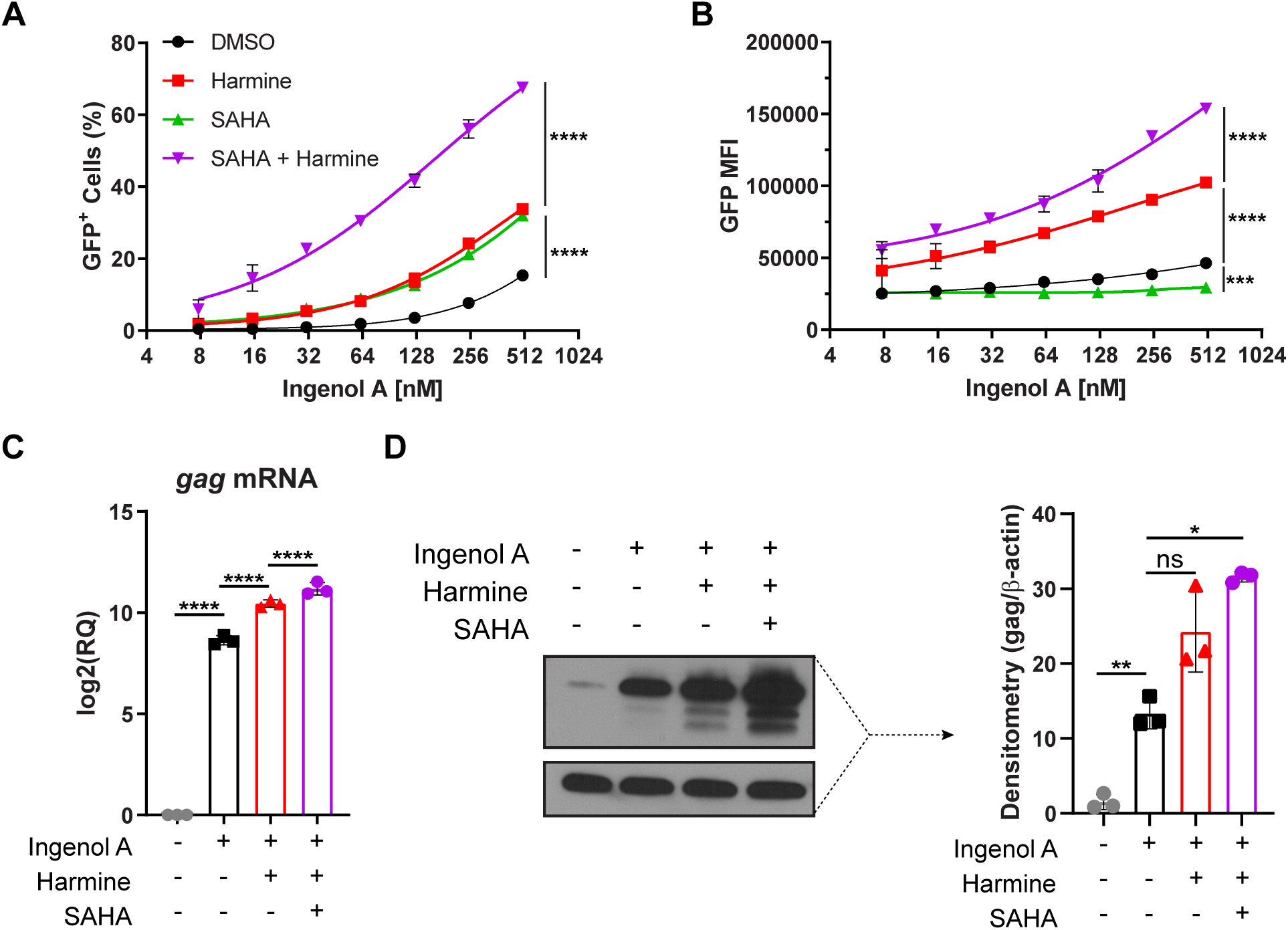
SAHA and harmine act synergistically to boost ingenol A activation of J-Lat cells. J-Lat 5A8 cells were pretreated with DMSO, harmine, SAHA, or SAHA + harmine for 30 minutes followed by ingenol A (31.25 nM) stimulation for 18 hours. **(A)** The percentage of GFP^+^ cells and **(B)** the mean fluorescence intensity (n = 3). Expression of *gag* mRNA and gag protein were measured by **(D)** qPCR and **(E)** western blot, respectively (n = 3). Statistical analysis was performed by calculating the area under the curve and performing a two-way-ANOVA corrected for multiple comparisons with Tukey’s test (A-B) or by one-way-ANOVA analysis corrected for multiple comparisons with Tukey’s test (C-D). *p<0.05; **p<0.01; ****p<0.0001; ns = not significant.

## Discussion

In the current study, we demonstrate that the plant-derived compound harmine can boost the efficacy of latent HIV-1 reactivation mediated by LRAs. Our data show that harmine has two effects: 1.) it increases the number of cells that are reactivated by LRAs, and 2.) it increases LTR promoter activity in reactivated cells (**Figure 7A-B**). Harmine has been shown to inhibit DYRK1A, a negative regulator of NFAT signaling ^17–19,24^. However, in the current study, we demonstrate that the boosting effect is independent of NFAT and DYRK1A. We found that harmine treatment results in reduced expression of HEXIM1, a negative regulator of transcription elongation mediated by pTEFb.

**Figure 7:**
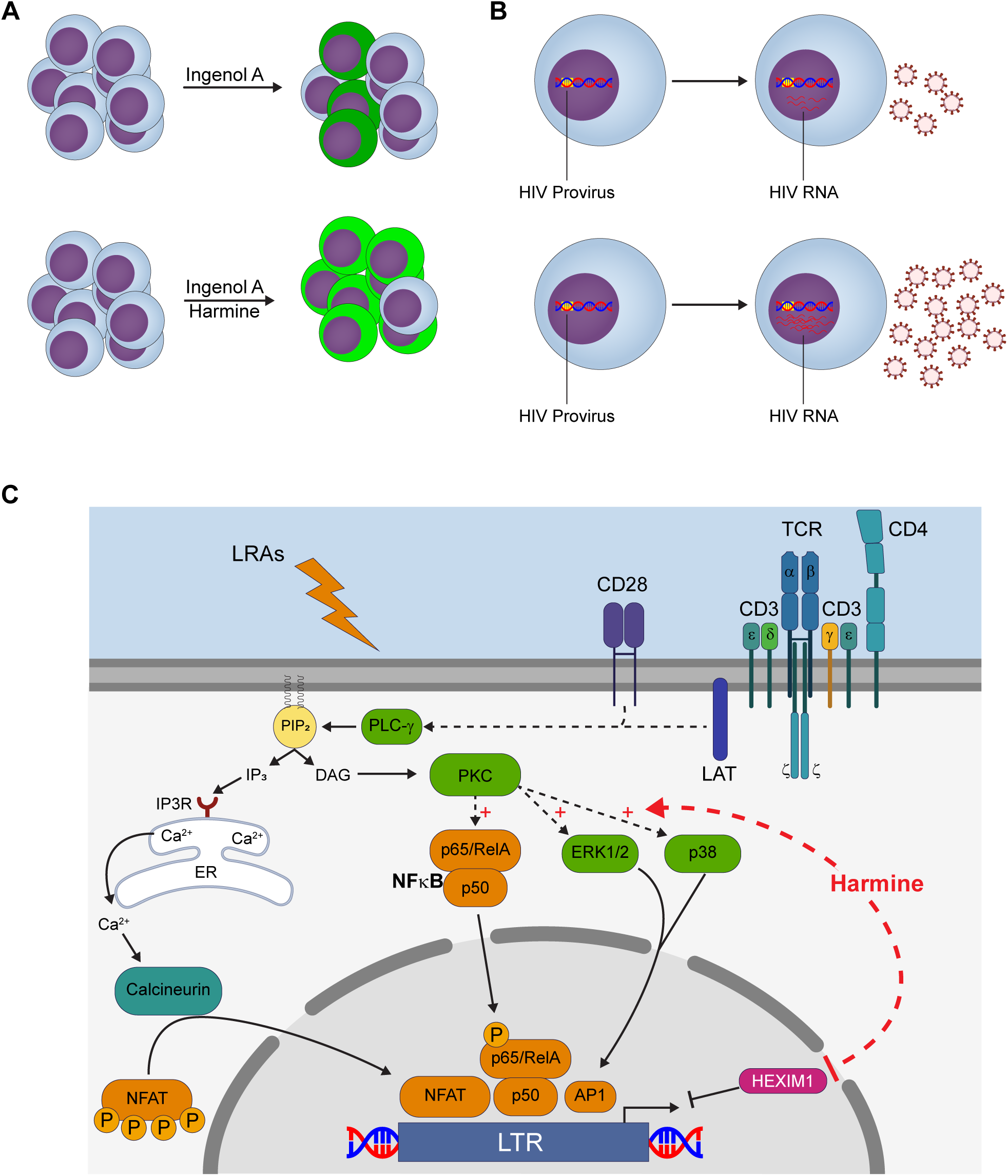
Harmine treatment results in increased HIV reactivation by PKC-agonists. **(A)** Combinatorial treatment of ingenol A and harmine results in increased frequency of GFP^+^ cells and increased MFI in GFP^+^ cells in J-Lat model. **(B)** Combinatorial treatment of ingenol A and harmine results in increased LTR activity and expression of HIV RNA. **(C)** Harmine treatment in combination with PKC agonists results in increased availability of transcription factor NFκB, increased MAPK p38 and ERK1/2 activity, and decreased HEXIM1 expression.

Previous studies have demonstrated that PKC agonists such as prostratin, bryostatin, and ingenol A can reactivate latent HIV-1 and that this mechanism is through NFκB activation ^26–33^. These studies have also demonstrated that reactivation of latent HIV-1 through PKC agonists is synergistic with latency reversal with HDAC inhibitors or JQ1 ^26–33^. This synergism is due to two different mechanisms for latency being targeted: availability of transcription factors and either epigenetic modifications or availability of pTEFb complexes. In our current study, we demonstrate that the compound harmine has an additive effect with PKC agonists and HDAC inhibitors.

A recent study by Booiman and colleagues demonstrated that the DYRK1A inhibitor, INDY, reactivated latently infected J-Lat cells without the use of a PKC agonist ^36^. In our hands, however, treatment of J-Lat cells with DYRK1A inhibitors alone was not sufficient to reactivate latently infected cells as measured by GFP expression. However, INDY treatment alone did result in increased *gag* mRNA expression (**Figure 1D**). The dose that we used for our study (20 µM) is much lower than the doses used by Booiman and colleagues. When higher doses of INDY were used in our studies, excessive toxicity for J-Lat cells was observed (not shown). Treatment with DYRK1A inhibitors only affected J-Lat cells that were stimulated with PKC agonists, SAHA, or TNF. The study by Booiman and colleagues used J-Lat clones 8.4 and A1, whereas we used clone 5A8, which may account for the differences ^36^.

We initially expected that harmine could enhance HIV-1 reactivation through inhibition of DYRK1A and increased availability of activated NFAT, but NFAT activity was not increased when using PKC agonists. Instead, we observed that NFκB activation was boosted by harmine. While it is not surprising that the mechanism of action is through NFκB for the PKC agonists used, it is surprising that harmine would enhance this pathway. The NFκB reporter cells utilize a luciferase-based system and therefore increased luciferase production does not necessarily mean increased NFκB phosphorylation *per se* and may just be a result of increased transcriptional elongation of luciferase transcripts. However, harmine did not affect ionomycin induced luciferase activity in the NFAT reporter cells suggesting that the increase in NFκB activity is not due to increased transcription of luciferase.

Since harmine is a bona fide DYRK1A inhibitor and has been shown to bind to DYRK1A in crystal structures, we hypothesized that harmine may be modifying the function of DYRK1A or directing it into a different pathway. Our data show that harmine treatment also boosts signaling through the MEK/ERK and p38 MAPK pathways when used in combination with a PKC agonist. This contrasted with INDY, which did not affect these pathways. While both harmine and INDY boosted the frequency of reactivated cells after treatment with PKC agonists, only harmine boosted the MFI of the reactivated cells. This suggested that harmine may have a secondary effect on DYRK1A. However, when we knocked out DYRK1A using CRISPR/Cas9, the boosting effect of harmine was still observed (**Figure 4**). This indicates that DYRK1A is not the target of harmine that is boosting HIV reactivation.

Microarray data demonstrated that harmine treatment reduced *HEXIM1* levels. This was confirmed at the protein level by western blot analysis (**Figure 5**). HEXIM is a component of the pTEFb complex. The pTEFb complex consists of multiple proteins including CycT1, CDK9, and HEXIM, and there are heterogeneous combinations of these proteins that determine the specific activity (or inactivity) of pTEFb. A 7SK snRNP complex containing HEXIM, CDK9, CycT1, MePCE, and LARP7 along with a 7SK snRNA holds CDK9 in an inactive complex ^37–40^. A key role for HIV Tat protein is to release CDK9/CycT1 pTEFb from 7SK snRNA so that it may drive HIV mRNA transcription ^41–43^. Alternatively, CDK9/CycT1 can be bound by BRD4 and this drives expression of host genes such as c-Myc ^44–47^. However, BRD4 competes with Tat for pTEFb ^48^. It has been shown that the BRD4 inhibitor JQ1 enhances Tat-induced HIV transcription and reversal of latency ^49,50^. Thus, the pTEFb transcription complex acts as a three-position switch relative to HIV and host gene mRNA expression. HEXIM-containing complexes are in the off position for all pTEFb-dependent genes. BRD4-containing complexes are in the off position for HIV transcription but the on position for host genes such as c-Myc. Finally, Tat-containing complexes are in the on position for HIV transcription.

HEXIM1, when part of the pTEFb complex, is inhibitory for HIV-1 replication ^35,39,40^ suggesting that the mechanism of boosting by harmine is through increasing the availability of active pTEFb complexes for recruitment by Tat. The microarray also revealed increased expression of CyclinT2. While CyclinT1 is the kinase normally associated with the pTEFb complex CyclinT2 has also been reported to associate with CDK9 in the PTEF-b complex. Unlike CyclinT1, which promotes Tat activity, CyclinT2 has been reported to be inhibitory for HIV-1 transcription ^51^.

In the J-Lat model, GFP acts as a surrogate measure for LTR activity. We also wanted to measure *gag* expression by Western blot and RT-qPCR to see if LTR promoter activity was increased. Ingenol A-induced *gag* expression was dramatically increased with the addition of harmine. Increased expression of viral proteins will also make the “kill” phase of the shock and kill approach more effective as it is necessary for targeting by immune cells. Signaling through the TCR activates NFκB and NFAT and it has been previously reported that TCR signaling enhances transcriptional elongation of latent HIV-1 by activating pTEFb through an ERK-dependent pathway ^52^. Interestingly, we found that treatment with ingenol A leads to ERK-1/2 phosphorylation and that this effect was enhanced by treatment with harmine (**Figure 3**). This may be the mechanism by which harmine treatment enhances LTR promoter activity through pTEFb.

Finally, we observed that harmine also boosted the effects of SAHA-mediated activation (**Supplemental Figure 3**). If harmine was mediating its boosting effect by the increased availability of transcription factors such as NFAT or NFκB, then we wouldn’t expect harmine to boost SAHA-mediated activation in the absence of PKC agonist stimulation. However, since we found that harmine treatment was downregulating *HEXIM1* expression (**Figure 5**), we tested to see if the combination of SAHA, harmine, and a PKC agonist would work additively to boost HIV reactivation. Indeed, we found that the combination treatment of the three was more potent than any combination of two reagents (**Figure 6**). This is likely due to harmine treatment increasing the availability of NFκB, enhancing the effect of the PKC agonist, while simultaneously boosting the activation of the MEK/ERK pathway which leads to increased transcript elongation. While SAHA inhibits HDACs and increases the availability of LTR promoters for transcription and with the increase in transcription by harmine this effect is also boosted.

Thus, harmine increases PKC-agonist-induced ERK1/2 signaling and results in downregulation of HEXIM1 which may result in increased transcriptional elongation through pTEFb. Furthermore, harmine enhances the effects of PKC agonist-induced HIV-1 reactivation through enhanced NFκB signaling and not through NFAT signaling (**Figure 7C**). We propose that harmine can be used in combination with existing clinically approved LRAs such as SAHA, bryostatin, or ingenol A to increase their efficacy *in vivo*.

## Methods

### Reagents and Resources

A list of key antibodies, inhibitors, commercial kits, and other reagents can be found in **Supplemental Table 2**.

### Reactivation Experiments

J-Lat 5A8 cells are treated with DMSO, harmine (20 µM), INDY (20 µM), or SAHA (5 µM) for 30 minutes followed by treatment with agonists for 18 hours unless otherwise indicated in figure legend.

### Cell Culture

J-Lat 5A8 cells, a kind gift from Warner Greene (University of California, San Francisco), have been previously described ^53^. Briefly, J-Lat 5A8 cells are Jurkat cells that are latently infected with a full-length provirus integrated into the *MAT2A* gene and have the *gfp* reporter gene in place of *nef*. A frameshift resulting in defective env production renders the cells non-infectious. Any stimulus that activates the LTR will result in transcription of *gfp*. J-Lat 5A8 cells and Jurkat luciferase reporter cells were cultured in RPMI 1640 with 2 mM L-glutamine (Corning) supplemented with 10% heat-inactivated FBS (Sigma-Aldrich, St. Louis, MO) and 100 U/mL penicillin-streptomycin.

### Primary CD4^+^ T Cells

Buffy coats were obtained from Life South Community Blood Center (Gainesville, FL) under approval by the Institutional Review Board at the University of Florida. PBMCs were processed as previously described ^54^ and CD4^+^ T cells were enriched with the CD4^+^ T Cell Isolation Kit (Miltenyi Biotec) according to the manufacturer’s instructions. CD4^+^ T cells were cultured in Advanced RPMI 1640 (Corning) supplemented with 2 mM GlutaMAX (Gibco), 10% heat-inactivated FBS (Sigma-Aldrich, St. Louis, MO), and 100 U/mL penicillin-streptomycin.

### Flow Cytometry and Flow Sorting

Cells were fixed in 2% paraformaldehyde and GFP fluorescence was measured using the BD Accuri™ C6 flow cytometer (BD Biosciences). Flow sorting was performed on unfixed cells the FACS Aria III (BD Biosciences) at the University of Florida Center for Immunology & Transplantation.

### AlamarBlue Assay

Jurkat T cells were treated with different doses of harmine, INDY, or an equivalent volume of DMSO overnight. Toxicity of the inhibitors was measured by adding 10 µL of AlamarBlue® Cell Viability Assay Reagent (Thermo Scientific) to 90 µL of cells and waiting for color development. Absorbance was measured at 570 and 600 nm. The percent reduction was calculated according to the manufacturer’s instructions.

### Western Blot

Western blots were performed as previously described ^54^.

### RT-qPCR

Total RNA was isolated with the RNeasy Plus Mini Kit (Qiagen). cDNA synthesis was carried out with High-Capacity cDNA Reverse Transcription Kit (Applied Biosystems, Foster City, CA) according to the manufacturer’s instructions. RT-qPCR reactions were carried out in SYBR™ Select Master Mix (Thermo Fisher Scientific, Waltham, MA) according to the manufacturer’s instructions. RT-qPCR reactions were performed on the StepOnePlus™ Real-Time PCR System (Applied Biosystems). Gag primers used were FWD: 5’-GAGCTAGAACGATTCGCAGTTA-3’; REV: 5’-CTGTCTGAAGGGATGGTTGTAG-3’.

### Luciferase Assays

Reporter cell lines were created by transducing Jurkat (E6.1) T cells with Cignal Lenti NFAT reporter (Cat# CLS-015L), Cignal Lenti NFκB reporter (Cat# CLS-013L), or Cignal Lenti Negative Control (CLS-NCL) lentiviral particles purchased from Qiagen. An equal volume of Bright-Glo™ Luciferase Assay System (Promega) was added to an equal volume of cells. After 5 minutes, the cells were lysed, and the lysate was transferred to a black Costar EIA/RIA polystyrene half area 96-well plate (Corning). Luminescence was measured with the VICTOR™ X4 Multi-Plate Reader (PerkinElmer).

### Microarrays

J-Lat 5A8 cells were treated with DMSO or harmine (20 µM) for 30 minutes. After pretreatment, the cells were treated with either medium or PMA (10 nM) for 2 hours. Total RNA was collected with the RNeasy Plus Mini Kit (Qiagen). Gene expression was assessed with GeneChip™ Human Transcriptome Array 2.0 arrays (Affymetrix) by the Interdisciplinary Center for Biotechnology Research at the University of Florida. The analysis was performed with Partek Genomics Suite v. 6.6 (Partek Inc., St. Louis, MO). CEL files were imported and the raw data were subjected to multi-array average (RMA) background correction and quantile normalization. Probesets were summarized by the median polish method and the summarized signals were transformed to log base 2. A one-way-ANOVA with contrast was performed to determine fold changes between PMA and PMA + harmine-treated groups. Transcripts that were significantly (step-up FDR < 0.05) upregulated or downregulated by at least 2-fold are listed in **Supplemental Table 1** (coding and noncoding) and **Table 1** (coding).

### Calcium Flux Assays

J-Lat 5A8 cells were incubated for 30 minutes with the Calcium Sensor Dye eFluor 514 (eBioscience) at a concentration of 2 µM with DMSO, harmine (20 µM), or INDY (20 µM). The cells were washed and resuspended in sample buffer (1X PBS, 0.1% BSA, 2 mM EDTA) containing DMSO, harmine (20 µM), or INDY (20 µM). Flow cytometry data were collected for one minute and then ionomycin (250 nM) or ingenol A (50 nM) was added and flow cytometry data were collected for an additional three minutes. The change in the area under the curve (δ AUC) was calculated by dividing the area under the curve after the addition of ionomycin or ingenol A divided by the area under the curve before the addition of ionomycin or ingenol A.

### CRISPR/Cas9 Knockout of DYRK1A

J-Lat 5A8 cells were co-transfected with two plasmids. One plasmid on the pCRISPR-CG01 backbone (GeneCopoeia) codes for recombinant Cas9 and a sgRNA (5’-GCCAAACATAAGTGACCAAC-3’) that targets exon 2 of DYRK1A. The second plasmid on the pDONOR-D01 backbone (GeneCopoeia) has a mCherry-T2A-Puro reporter cassette flanked by homology regions adjacent to the sgRNA target site in the genome. After co-transfection, the J-Lat cells were selected with puromycin (1 µg/mL).

### Statistical Analysis

All statistical analyses were performed using GraphPad Prism for Windows (GraphPad Software, La Jolla, California). A p-value of less than 0.05 was considered statistically significant.

## Data Availability

Microarray data are available in Gene Expression Omnibus (GEO) under accession GSE136172.

## Acknowledgments

The authors would like to thank Warner Greene (University of California, San Francisco) for generously providing the J-Lat 5A8 cells used in the study. This work was supported by funding from the National Institute of Allergy and Infectious Diseases grants R56AI108434 and R56AI122813. J.P.T. was supported by the Ruth L. Kirschstein National Research Service Award Institutional Research Training Grant T32AI007110.

## Author Contributions

J.P.T. helped conceive of the project, designed and performed experiments, analyzed data, took the lead in writing the manuscript, and contributed to the interpretation of the results. L.H.A. performed the calcium flux and luciferase experiments. D.L.A. performed CRISPR transfections to generate the DYRK1A^-/-^ knockout cell line. M.N.C. created the Jurkat T cell NFκB and NFAT reporter cell lines. M.A.W. designed and directed the project, performed experiments, analyzed data, and contributed to the interpretation of the results.

## Figure Legends

**Supplemental Figure 1. Dose titration of DYRK1A inhibitors.**

**(A)** Chemical structures for harmine and INDY were generated with BIOVIA Draw v18.1 (Dassault Systèmes). **(B)** J-Lat 5A8 cells were pretreated with harmine or INDY for 30 minutes followed by activation with PMA (1.25 nM) for 18 hours (n = 1). **(C)** CD4^+^ T cells were pretreated with harmine or INDY followed by stimulation with 1X Cell Stimulation Cocktail (81 nM PMA; 1.34 µM ionomycin) overnight. Supernatants were collected and TNF was measured by ELISA (n = 1). **(D)** Viability was measured by AlamarBlue Assay. Decreased reduction capacity indicates cellular toxicity (n = 1).

**Supplemental Figure 2. DYRK1A inhibitors enhance the efficacy of other PKC agonists.**

J-Lat 5A8 cells were reactivated by titrating ingenol A, PMA, or TNF in the presence of inhibitors. The percentage of GFP^+^ cells and the geometric mean fluorescence intensity were measured by flow cytometry (n = 3).

**Supplemental Figure 3. SAHA has no effect on the MFI of GFP**^**+**^ **cells but does boost the percentage of reactivated cells.**

SAHA was titrated on J-Lat 5A8 cells pretreated with DMSO, harmine, or INDY for 30 minutes. The percentage of GFP^+^ cells and the mean fluorescence intensity of GFP^+^ cells was measured by flow cytometry (n = 1).

## References

1. Siliciano, R. F. & Greene, W. C. HIV latency. Cold Spring Harb. Perspect. Med. 1, a007096 (2011).

2. Folks, T. et al. Induction of HTLV-III/LAV from a nonvirus-producing T-cell line: implications for latency. Science 231, 600–2 (1986).

3. Davey, R. T. et al. HIV-1 and T cell dynamics after interruption of highly active antiretroviral therapy (HAART) in patients with a history of sustained viral suppression. Proc. Natl. Acad. Sci. U. S. A. 96, 15109–14 (1999).

4. Hamer, D. H. Can HIV be Cured? Mechanisms of HIV persistence and strategies to combat it. Curr. HIV Res. 2, 99–111 (2004).

5. Deeks, S. G. HIV: Shock and kill. Nature 487, 439–40 (2012).

6. Archin, N. M. et al. Expression of latent human immunodeficiency type 1 is induced by novel and selective histone deacetylase inhibitors. AIDS 23, 1799–806 (2009).

7. Archin, N. M. et al. Expression of latent HIV induced by the potent HDAC inhibitor suberoylanilide hydroxamic acid. AIDS Res. Hum. Retroviruses 25, 207–12 (2009).

8. Contreras, X. et al. Suberoylanilide hydroxamic acid reactivates HIV from latently infected cells. J. Biol. Chem. 284, 6782–9 (2009).

9. Lehrman, G. et al. Depletion of latent HIV-1 infection in vivo: a proof-of-concept study. Lancet (London, England) 366, 549–55 (2005).

10. Archin, N. M. et al. Administration of vorinostat disrupts HIV-1 latency in patients on antiretroviral therapy. Nature 487, 482–5 (2012).

11. Zhu, Y. et al. Transcription elongation factor P-TEFb is required for HIV-1 tat transactivation in vitro. Genes Dev. 11, 2622–32 (1997).

12. Mancebo, H. S. et al. P-TEFb kinase is required for HIV Tat transcriptional activation in vivo and in vitro. Genes Dev. 11, 2633–44 (1997).

13. Macian, F. NFAT proteins: key regulators of T-cell development and function. Nat. Rev. Immunol. 5, 472–84 (2005).

14. Gwack, Y. et al. A genome-wide Drosophila RNAi screen identifies DYRK-family kinases as regulators of NFAT. Nature 441, 646–50 (2006).

15. Arron, J. R. et al. NFAT dysregulation by increased dosage of DSCR1 and DYRK1A on chromosome 21. Nature 441, 595–600 (2006).

16. Patel, K., Gadewar, M., Tripathi, R., Prasad, S. K. & Patel, D. K. A review on medicinal importance, pharmacological activity and bioanalytical aspects of beta-carboline alkaloid ‘“Harmine”’. Asian Pac. J. Trop. Biomed. 2, 660–4 (2012).

17. Adayev, T., Wegiel, J. & Hwang, Y.-W. Harmine is an ATP-competitive inhibitor for dual-specificity tyrosine phosphorylation-regulated kinase 1A (Dyrk1A). Arch. Biochem. Biophys. 507, 212–8 (2011).

18. Bain, J. et al. The selectivity of protein kinase inhibitors: a further update. Biochem. J. 408, 297–315 (2007).

19. Göckler, N. et al. Harmine specifically inhibits protein kinase DYRK1A and interferes with neurite formation. FEBS J. 276, 6324–37 (2009).

20. Slotkin, T. & DiStefano, V. Urinary metabolities of harmine in the rat and their inhibition of monoamine oxidase. Biochem. Pharmacol. 19, 125–31 (1970).

21. Blum, B. et al. Harmine antagonism of drug-induced extra-pyramidal disturbances. Psychopharmacologia 6, 307–10 (1964).

22. Tung, Y. C., Hwang, Y. S. & Wu, C. Y. STUDIES ON BETA-CARBOLINES. 3. EFFECT OF HARMAN HYDROCHLORIDE ON MONOAMINE OXIDASE AND THE UTERINE CONTRACTILITY INDUCED BY N-MONOMETHYLTRYPTAMINE HYDROCHLORIDE. Tsa. Chih. Gaoxiong Yi Xue Yuan. Tong Xue Hui 64, 44–50 (1965).

23. Yasuhara, H. Studies on monoamine oxidase (report XXIV). Effect of harmine on monoamine oxidase. Jpn. J. Pharmacol. 24, 523–33 (1974).

24. Egusa, H. et al. The small molecule harmine regulates NFATc1 and Id2 expression in osteoclast progenitor cells. Bone 49, 264–74 (2011).

25. Ogawa, Y. et al. Development of a novel selective inhibitor of the Down syndrome-related kinase Dyrk1A. Nat. Commun. 1, 86 (2010).

26. Williams, S. A. et al. Prostratin antagonizes HIV latency by activating NF-kappaB. J. Biol. Chem. 279, 42008–17 (2004).

27. Laird, G. M. et al. Ex vivo analysis identifies effective HIV-1 latency-reversing drug combinations. J. Clin. Invest. 125, 1901–12 (2015).

28. Reuse, S. et al. Synergistic activation of HIV-1 expression by deacetylase inhibitors and prostratin: implications for treatment of latent infection. PLoS One 4, e6093 (2009).

29. Perez, M. et al. Bryostatin-1 Synergizes with Histone Deacetylase Inhibitors to Reactivate HIV-1 from Latency. Curr. HIV Res. 8, 418–429 (2010).

30. Martínez-Bonet, M. et al. Synergistic Activation of Latent HIV-1 Expression by Novel Histone Deacetylase Inhibitors and Bryostatin-1. Sci. Rep. 5, 16445 (2015).

31. Jiang, G. et al. Synergistic Reactivation of Latent HIV Expression by Ingenol-3-Angelate, PEP005, Targeted NF-kB Signaling in Combination with JQ1 Induced p-TEFb Activation. PLoS Pathog. 11, e1005066 (2015).

32. Brogdon, J., Ziani, W., Wang, X., Veazey, R. S. & Xu, H. In vitro effects of the smallmolecule protein kinase C agonists on HIV latency reactivation. Sci. Rep. 6, 39032 (2016).

33. Díaz, L. et al. Bryostatin activates HIV-1 latent expression in human astrocytes through a PKC and NF-κB-dependent mechanism. Sci. Rep. 5, 12442 (2015).

34. Chen, F. X., Smith, E. R. & Shilatifard, A. Born to run: Control of transcription elongation by RNA polymerase II. Nat. Rev. Mol. Cell Biol. 19, 464–478 (2018).

35. Barboric, M. et al. Tat competes with HEXIM1 to increase the active pool of P-TEFb for HIV-1 transcription. Nucleic Acids Res. 35, 2003–12 (2007).

36. Booiman, T., Loukachov, V. V., van Dort, K. A., van’t Wout, A. B. & Kootstra, N. A. DYRK1A Controls HIV-1 Replication at a Transcriptional Level in an NFAT Dependent Manner. PLoS One 10, e0144229 (2015).

37. Nguyen, V. T., Kiss, T., Michels, A. A. & Bensaude, O. 7SK small nuclear RNA binds to and inhibits the activity of CDK9/cyclin T complexes. Nature 414, 322–5 (2001).

38. Yang, Z., Zhu, Q., Luo, K. & Zhou, Q. The 7SK small nuclear RNA inhibits the CDK9/cyclin T1 kinase to control transcription. Nature 414, 317–22 (2001).

39. Michels, A. a et al. Binding of the 7SK snRNA turns the HEXIM1 protein into a P-TEFb (CDK9/cyclin T) inhibitor. EMBO J. 23, 2608–19 (2004).

40. Yik, J. H. N. et al. Inhibition of P-TEFb (CDK9/Cyclin T) kinase and RNA polymerase II transcription by the coordinated actions of HEXIM1 and 7SK snRNA. Mol. Cell 12, 971–82 (2003).

41. Fujinaga, K. et al. Dynamics of human immunodeficiency virus transcription: P-TEFb phosphorylates RD and dissociates negative effectors from the transactivation response element. Mol. Cell. Biol. 24, 787–95 (2004).

42. Ivanov, D., Kwak, Y. T., Guo, J. & Gaynor, R. B. Domains in the SPT5 protein that modulate its transcriptional regulatory properties. Mol. Cell. Biol. 20, 2970–83 (2000).

43. Ping, Y. H. & Rana, T. M. DSIF and NELF interact with RNA polymerase II elongation complex and HIV-1 Tat stimulates P-TEFb-mediated phosphorylation of RNA polymerase II and DSIF during transcription elongation. J. Biol. Chem. 276, 12951–8 (2001).

44. Yang, Z. et al. Recruitment of P-TEFb for stimulation of transcriptional elongation by the bromodomain protein Brd4. Mol. Cell 19, 535–45 (2005).

45. Jang, M. K. et al. The bromodomain protein Brd4 is a positive regulatory component of P-TEFb and stimulates RNA polymerase II-dependent transcription. Mol. Cell 19, 523–34 (2005).

46. Yang, Z., He, N. & Zhou, Q. Brd4 recruits P-TEFb to chromosomes at late mitosis to promote G1 gene expression and cell cycle progression. Mol. Cell. Biol. 28, 967–76 (2008).

47. Dey, A., Nishiyama, A., Karpova, T., McNally, J. & Ozato, K. Brd4 marks select genes on mitotic chromatin and directs postmitotic transcription. Mol. Biol. Cell 20, 4899–909 (2009).

48. Li, Z., Guo, J., Wu, Y. & Zhou, Q. The BET bromodomain inhibitor JQ1 activates HIV latency through antagonizing Brd4 inhibition of Tat-transactivation. Nucleic Acids Res. 41, 277–87 (2013).

49. Banerjee, C. et al. BET bromodomain inhibition as a novel strategy for reactivation of HIV-1. J. Leukoc. Biol. 92, 1147–54 (2012).

50. Bartholomeeusen, K., Xiang, Y., Fujinaga, K. & Peterlin, B. M. Bromodomain and extra-terminal (BET) bromodomain inhibition activate transcription via transient release of positive transcription elongation factor b (P-TEFb) from 7SK small nuclear ribonucleoprotein. J. Biol. Chem. 287, 36609–16 (2012).

51. Napolitano, G. et al. The CDK9-associated cyclins T1 and T2 exert opposite effects on HIV-1 Tat activity. AIDS 13, 1453–9 (1999).

52. Kim, Y. K., Mbonye, U., Hokello, J. & Karn, J. T-Cell receptor signaling enhances transcriptional elongation from latent HIV proviruses by activating P-TEFb through an ERK-dependent pathway. J. Mol. Biol. 410, 896–916 (2011).

53. Chan, J. K., Bhattacharyya, D., Lassen, K. G., Ruelas, D. & Greene, W. C. Calcium/calcineurin synergizes with prostratin to promote NF-κB dependent activation of latent HIV. PLoS One 8, e77749 (2013).

54. Taylor, J. P. et al. CRISPR/Cas9 knockout of USP18 enhances type I IFN responsiveness and restricts HIV-1 infection in macrophages. J. Leukoc. Biol. 1–16 (2018). doi:10.1002/JLB.3MIA0917-352R

