## Supplemental Files for "Harmine enhances the activity of the HIV-1 latency-reversing agents ingenol A and SAHA"

### Supplemental Figure 1

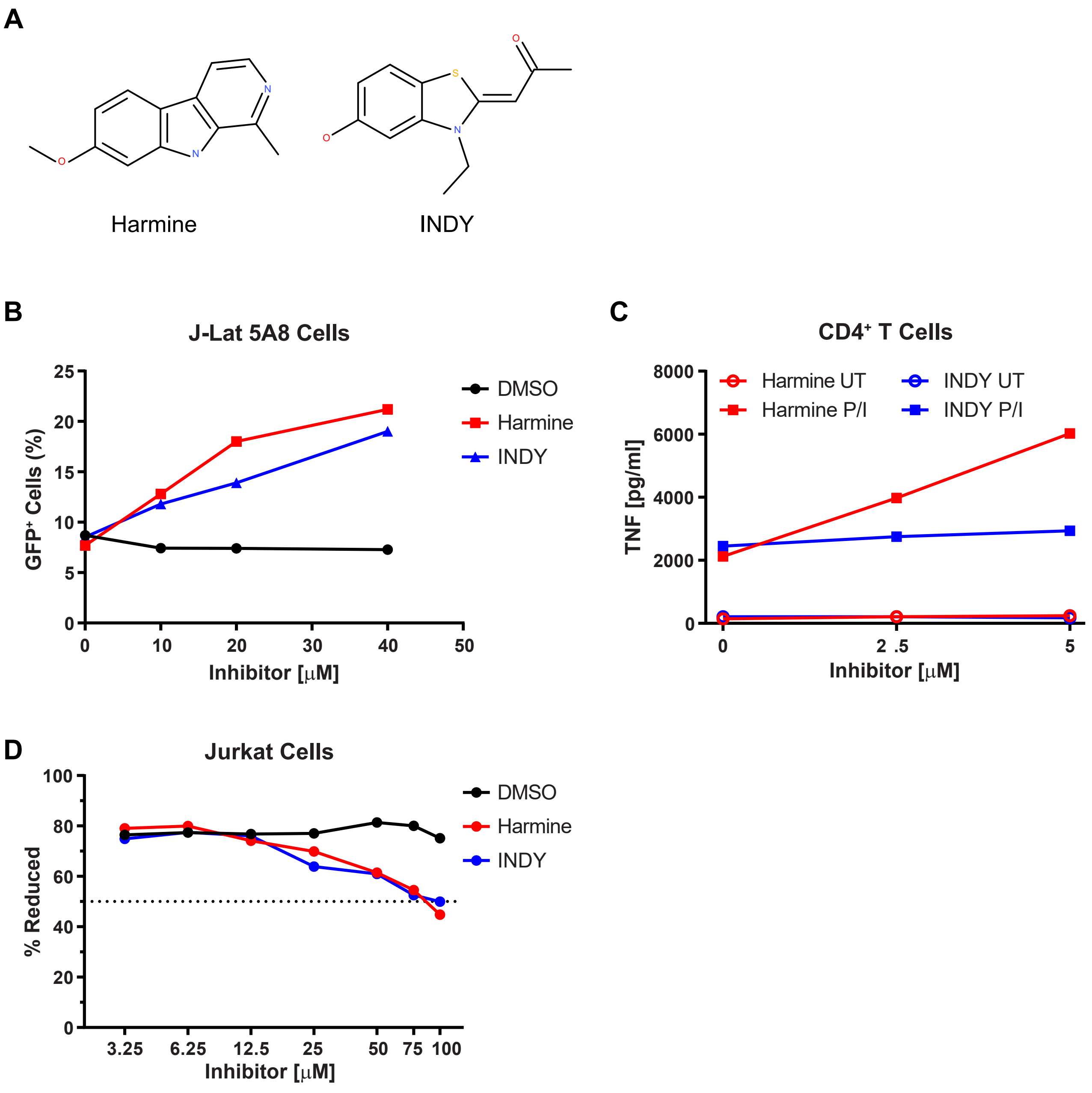

### Supplemental Figure 2

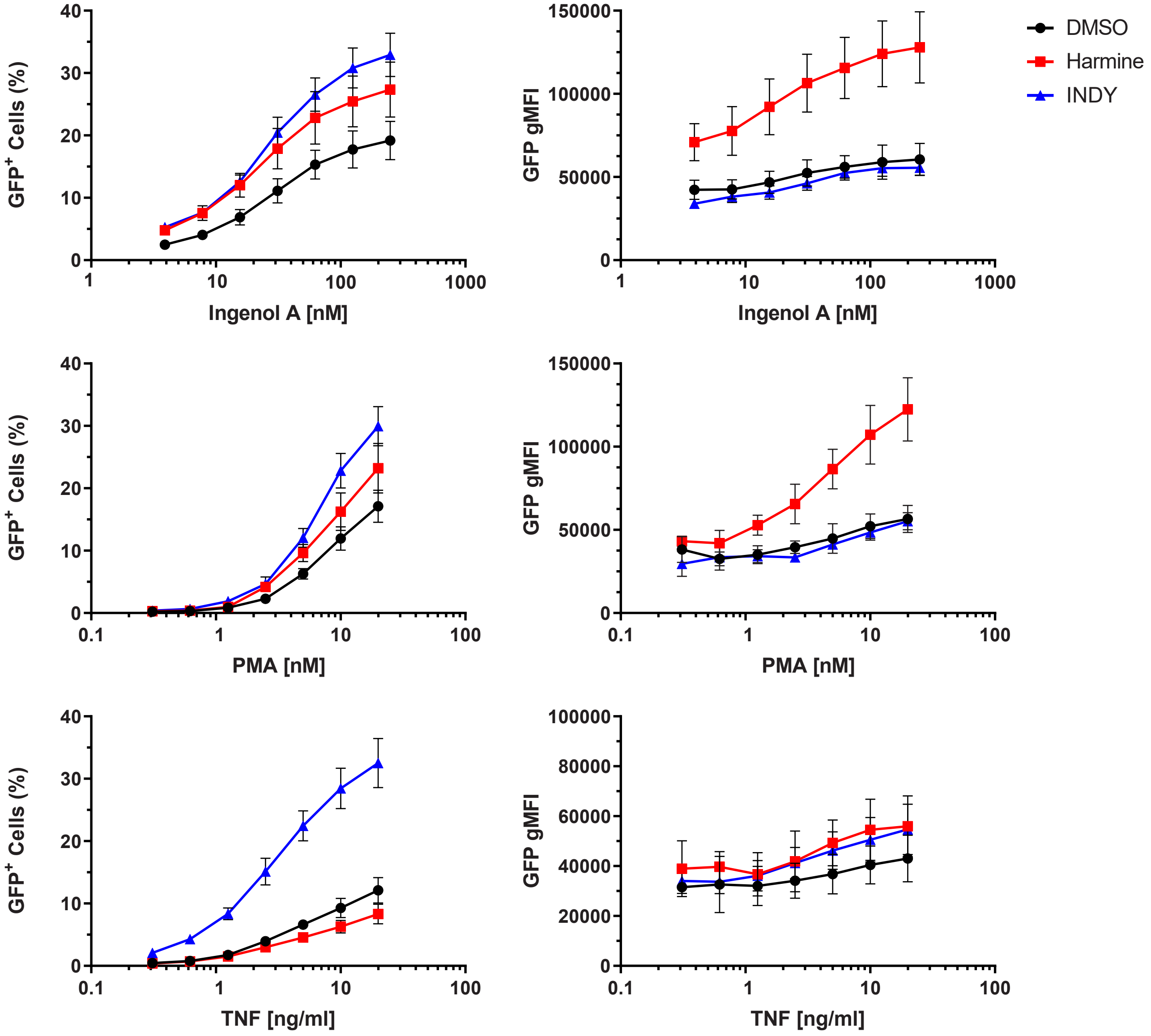

### Supplemental Figure 3

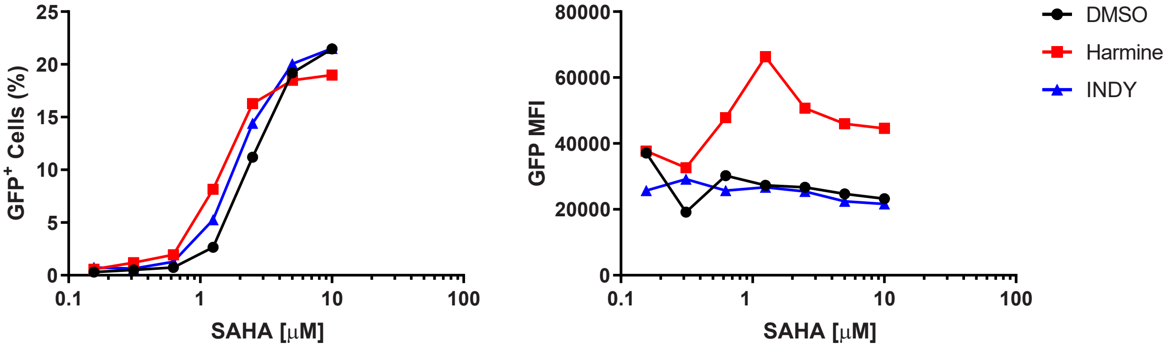

**Supplemental Table 1. Transcripts significantly upregulated with PMA + harmine vs PMA treatment**

| Transcript ID | Gene Symbol | RefSeq | p-value | Fold-Change |
|---|---|---|---|---|
| TC17000580.hg.1 | HEXIM1 | NM_006460 | 1.50E-08 | -3.29431 |
| TC01005596.hg.1 | | --- | 0.00012626 | -2.73307 |
| TC15000300.hg.1 | CHAC1 | NM_001142776 | 8.53E-08 | -2.52968 |
| TC01005769.hg.1 | | --- | 0.000141118 | -2.49202 |
| TC0X000277.hg.1 | USP27X | NM_001145073[a] | 1.36E-05 | -2.44667 |
| TC03002656.hg.1 | SKIL | BC041913 | 0.00013883 | -2.4415 |
| TC02004776.hg.1 | | --- | 0.000241331 | -2.43804 |
| TC02003463.hg.1 | | --- | 3.56E-05 | -2.43533 |
| TC21000594.hg.1 | | --- | 5.22E-05 | -2.42017 |
| TC01000837.hg.1 | LRRC8B | NM_001134476 | 2.42E-07 | -2.3876 |
| TC04001570.hg.1 | SLC7A11 | NM_014331 | 1.02E-05 | -2.37159 |
| 47423604_st | | --- | 0.000545199 | -2.3214 |
| TC03002567.hg.1 | RASA2 | AK025305 | 0.000178273 | -2.3009 |
| TC09002259.hg.1 | | --- | 0.000329717 | -2.28592 |
| TC14001777.hg.1 | | --- | 4.12E-05 | -2.23246 |
| TC01005963.hg.1 | | --- | 1.56E-05 | -2.23177 |
| TC19000250.hg.1 | IL27RA | NM_004843 | 2.71E-07 | -2.1761 |
| TC01005597.hg.1 | | --- | 2.22E-07 | -2.14946 |
| TC0X001705.hg.1 | USP27X | AY672104[a] | 1.46E-05 | -2.13768 |
| TC04001530.hg.1 | ANKRD50 | NM_001167882 | 3.69E-07 | -2.12436 |
| TC05003082.hg.1 | | --- | 7.27E-05 | -2.11168 |
| TC01003710.hg.1 | | --- | 0.000130796 | -2.06941 |
| TC06001830.hg.1 | RAB23 | NM_016277 | 4.59E-05 | -2.03377 |
| TC12002394.hg.1 | | --- | 0.000456661 | -2.00521 |
| TC09001204.hg.1 | TMEM2 | NM_001135820 | 1.88E-07 | -2.0027 |
| TC05001987.hg.1 | SOX30 | NM_007017 | 1.37E-07 | 2.10846 |
| TC02003598.hg.1 | CCNT2 | NR_037649 | 2.86E-06 | 2.2142 |
| TC02003784.hg.1 | | --- | 1.06E-05 | 2.23854 |
| TC17002650.hg.1 | | --- | 3.99E-06 | 2.26835 |
| TC17002649.hg.1 | | --- | 4.80E-05 | 2.35203 |
| JUC_ASC002193 | | | 7.03E-05 | 2.35559 |
| TC14001996.hg.1 | | --- | 7.08E-05 | 2.61315 |
| TC03002295.hg.1 | | --- | 3.16E-07 | 2.9908 |
| TC07001959.hg.1 | | --- | 2.99E-07 | 3.01515 |
| TC04000750.hg.1 | | --- | 7.63E-07 | 3.75633 |

[a] Probes bind to the same transcript. Only NM_001145073 is included in heat map in figure 5.

**Supplemental Table 2. Key reagents and resources**

| REAGENT or RESOURCE | SOURCE | IDENTIFIER |
|---|---|---|
| **Experimental Models: Cell Lines** | | |
| J-Lat 5A8 Cells | Warner Greene, University of California San Francisco | N/A |
| Jurkat (E6.1) Cells | ATCC | Cat# TIB-152; RRID:CVCL_0367 |
| **Antibodies** | | |
| Anti-HIV-1 p24 | Abcam | Cat# ab9071, RRID:AB_306981 |
| Anti-HSP90 | Cell Signaling | Cat# 4877, RRID:AB_2233307 |
| Anti-phospho-p44/42 MAPK | Cell Signaling | Cat# 5726, RRID:AB_2797617 |
| Anti-phospho-AKT | Cell Signaling | Cat# 2965, RRID:AB_2255933 |
| Anti-DYRK1A | Cell Signaling | Cat# 8765, RRID:AB_2797660 |
| Anti-phospho-p38 MAPK | Cell Signaling | Cat# 4511, RRID:AB_2139682 |
| Anti-HEXIM1 | Cell Signaling | Cat# 12604, RRID:AB_2797969 |
| Anti-β-actin | Cell Signaling | Cat# 4967, RRID:AB_330288 |
| **Lentivirus** | | |
| Cignal Lenti Negative Control (luc) | Qiagen | Cat# CLS-NCL |
| Cignal Lenti NFκB Reporter (luc) | Qiagen | Cat# CLS-013L |
| Cignal Lenti NFAT Reporter (luc) | Qiagen | Cat# CLS-015L |
| **Inhibitors** | | |
| U0126 | Cell Signaling | Cat# 9903 |
| Harmine | Tocris | Cat# 5075 |
| INDY | Tocris | Cat# 4997 |
| IKK 16 | MedChem Express | Cat# HY-13687 |
| **Latency Reactivating Agents** | | |
| SAHA (Vorinostat) | NIH AIDS Reagents Program | Cat# 12130 |
| Ionomycin | Cell Signaling | Cat# 9995 |
| Ingenol A (PEP005) | Tocris | Cat# 4054 |
| PMA | Sigma | Cat# P1585 |
| Recombinant Human TNF | PeproTech | Cat# 300-01A |
| **Commercial Kits** | | |
| RNeasy Plus Mini Kit | Qiagen | Cat# 74136 |
| High Capacity cDNA Reverse Transcription Kit | Applied Biosystems | Cat# 4368814 |
| SYBR Select Master Mix | Thermo Fisher | Cat# 4472918 |
| Calcium Sensor Dye eFluor 514 | eBioscience | Cat# 65-0859 |

| | | |
|---|---|---|
| Bright-Glo Luciferase Assay System | Promega | Cat# E2620 |
| BD OptEIA Human TNF ELISA Set | BD Biosciences | Cat# 555212 |
| AlamarBlue Cell Viability Assay Reagent | Thermo Fisher | Cat# 88951 |
| Cell Stimulation Cocktail (500X) | eBioscience | Cat# 00-4970 |
| $CD4^+$ T Cell Isolation Kit, Human | Miltenyi Biotec | Cat# 130-096-533 |
| **Deposited Data** | | |
| Microarray Data | This Paper | GEO# GSE136172 |
